## Supplement 1 for "Testing hypotheses of marsupial brain size variation using phylogenetic multiple imputations and a Bayesian comparative framework"

### Supplementary material

Table with data sources

| Trait | Units | Rationale | Reference |
| --- | --- | --- | --- |
| Brain | mm3 |  | (Weisbecker et al., 2015) |
| Body | grams |  | (Birdlife International, 2016; Flannery, 2013; Myers et al., 2006; van Dyck, Gynther, & Baker, 2013; Weisbecker, Ashwell, & Fisher, 2013) |
| Origin | 1 – Australia, 2 – New Guinea, 3 – Americas | Different origins predispose different influence of seasonality, predation pressure, food abundance. | (Flannery, 2013; Myers et al., 2006; van Dyck et al., 2013) |
| Status | 1 - Common, abundant, 2 - Vulnerable, endangered, rare, declining, limited  3 - Extinct | Highly threatened mammals are known to have larger relative brain sizes (Abelson, 2016) | (Birdlife International, 2016; van Dyck et al., 2013) |
| Diurnality | 1- Nocturnal, 2 – Diurnal, 3 - Crepuscular or not fully nocturnal | Nocturnal animals are considered larger brained, but daily activity is related to more complex predator avoidance techniques. | (Flannery, 2013; van Dyck et al., 2013; Weisbecker et al., 2015) |
| Arboreality | 1 - Arboreal or scansorial, 2 - Terrestrial | Arboreal environment is considered more cognitively demanding. | (Flannery, 2013; van Dyck et al., 2013; Weisbecker et al., 2015) |
| Shelter safety | 1 - Protected (burrow/nest in a tree hollow), 2 - Intermediate (tree canopy/hollow log/under rock/nest on the ground or in a soil crack), 3 - Open (under shrubs/in grass/tree shade) | Proxy for predation as selection pressure for larger brains. (Reddon, Chouinard-Thuly, Leris, & Reader, 2018) | (Flannery, 2013; van Dyck et al., 2013; (Weisbecker et al., 2015) |
| Diet | 1 - >50% grass/browse, 2 - Seeds, grass, roots, leaves, fruit, invertebrates, 3 - Nectar, fruit, invertebrates, 4 - >50% invertebrate/vertebrate | Foraging complexity and diet rich in nutrients have been shown to influence brain size | (Flannery, 2013; van Dyck et al., 2013; Weisbecker et al., 2015) |
| Group living | 1 – No, 2 - Yes | Measure of social complexity, which imposes greater interaction and recognition demands | (Flannery, 2013; Myers et al., 2006; van Dyck et al., 2013; Weisbecker et al., 2015) |
| Parental care | 1 – No, 2 - Yes | Parental investment is known to positively influence brain size (Isler & van Schaik, 2012) | (Flannery, 2013; Myers et al., 2006; van Dyck et al., 2013; Weisbecker et al., 2015) |
| Mating system | 1 – Promiscuous, 2 - Complex (polygamous/monogamous) | Complex mating systems require more cognitive complexity and usually result in higher parental investment (Schillaci, 2006) | (Flannery, 2013; van Dyck et al., 2013; Weisbecker et al., 2015) |
| Litter size | Average litter per reproductive episode | Constraint on maternal investment. | (van Dyck et al., 2013; Weisbecker et al., 2015) |
| Weaning age | Months | Constraint on maternal investment. | (Weisbecker et al., 2015) |
| Home range | Hectares | Larger home ranges usually imply increased cognitive complexity related to orientation (Clutton‐Brock & Harvey, 1980) | (Myers et al., 2006; van Dyck et al., 2013; Weisbecker et al., 2015) |
| Population density | Individuals per hectare | Increased population density is a proxy of increased interaction and social tolerance. | (van Dyck et al., 2013; (Weisbecker et al., 2015) |
| FMR | Field metabolic rate | Measure of metabolic turnover in the wild. | (Riek & Bruggeman, 2013) |
| Torpor | 0 – No, 1 – Yes | Torporing has been shown to be costly to the maintenance of large brains (Heldstab, Isler, & van Schaik, 2018) | (Geiser & Körtner, 2010; McNab, 2008; Ruf & Geiser, 2015) |
| Play | 1 – No, 2 – Rudimentary, 3 - Complex | Proxy for cognitive ability. Play has been shown to correlate with larger brains in birds and mammals (Iwaniuk, Nelson, & Pellis, 2001) | (Ashwell, 2008; Iwaniuk et al., 2001) |

Evolutionary mode analysis

| Origin | Brain | Body |
| --- | --- | --- |
| Australia | EB  95.3% | EB  69.5% |
| New Guinea | EB  53.8 % | BM  60.3% |
| Americas | BM  60.1% | BM  60.9% |

Table detailing the most probable models of continuous character evolution for brain and body size respectively, for marsupials from three different geographic regions and the AIC loading based on comparison between BM (Brownian motion), OU (Ornstein-Uhlenbeck), and EB (Early burst) models.

Dataset

Imputed datasets

R Code

Graphs

Heldstab, S. A., Isler, K., & van Schaik, C. P. (2018). Hibernation constrains brain size evolution in mammals. *J Evol Biol, 31*(10), 1582-1588. doi:10.1111/jeb.13353

Iwaniuk, A. N., Nelson, J. E., & Pellis, S. M. (2001). Do big-brained animals play more? Comparative analyses of play and relative brain size in mammals. *Journal of Comparative Psychology, 115*(1), 29-41. doi:10.1037/0735-7036.115.1.29
