## Supplement 2 for "Testing hypotheses of marsupial brain size variation using phylogenetic multiple imputations and a Bayesian comparative framework"

MCMCglmm models

After evaluating VIFs, some models were run with interaction term and some without, as follows:

formula_dev <- Brain ~ Weaning.age + Litter.size + BodyN

formula_soc <- Brain ~ Group.living + Parental.care + Mating.system + Population.density + BodyN

formula_env <- Brain ~ DiurnalityN + Shelter.safety + Arboreality + Diet + HR + BodyN

formula_ori <- Brain ~ Origin * BodyN

formula_vul <- Brain ~ Status * BodyN

formula_tor <- Brain ~ Torpor * BodyN

formula_pla <- Brain ~ Play * BodyN

formula_fmr <- Brain ~ FMR.Riek * BodyN

Missingness analysis

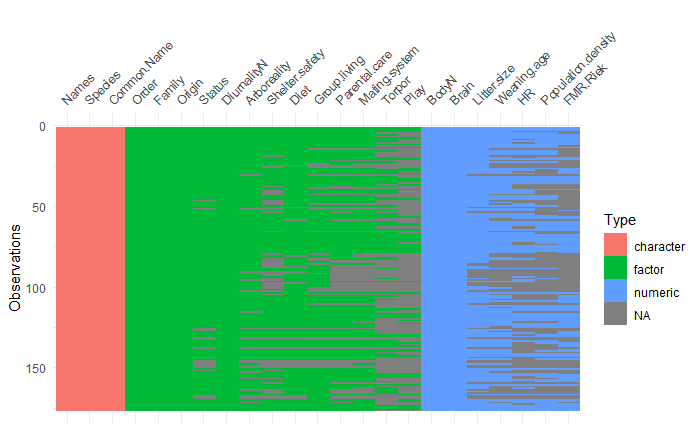

Figure 1a. Pattern of missingness in all variables divided by type of variable
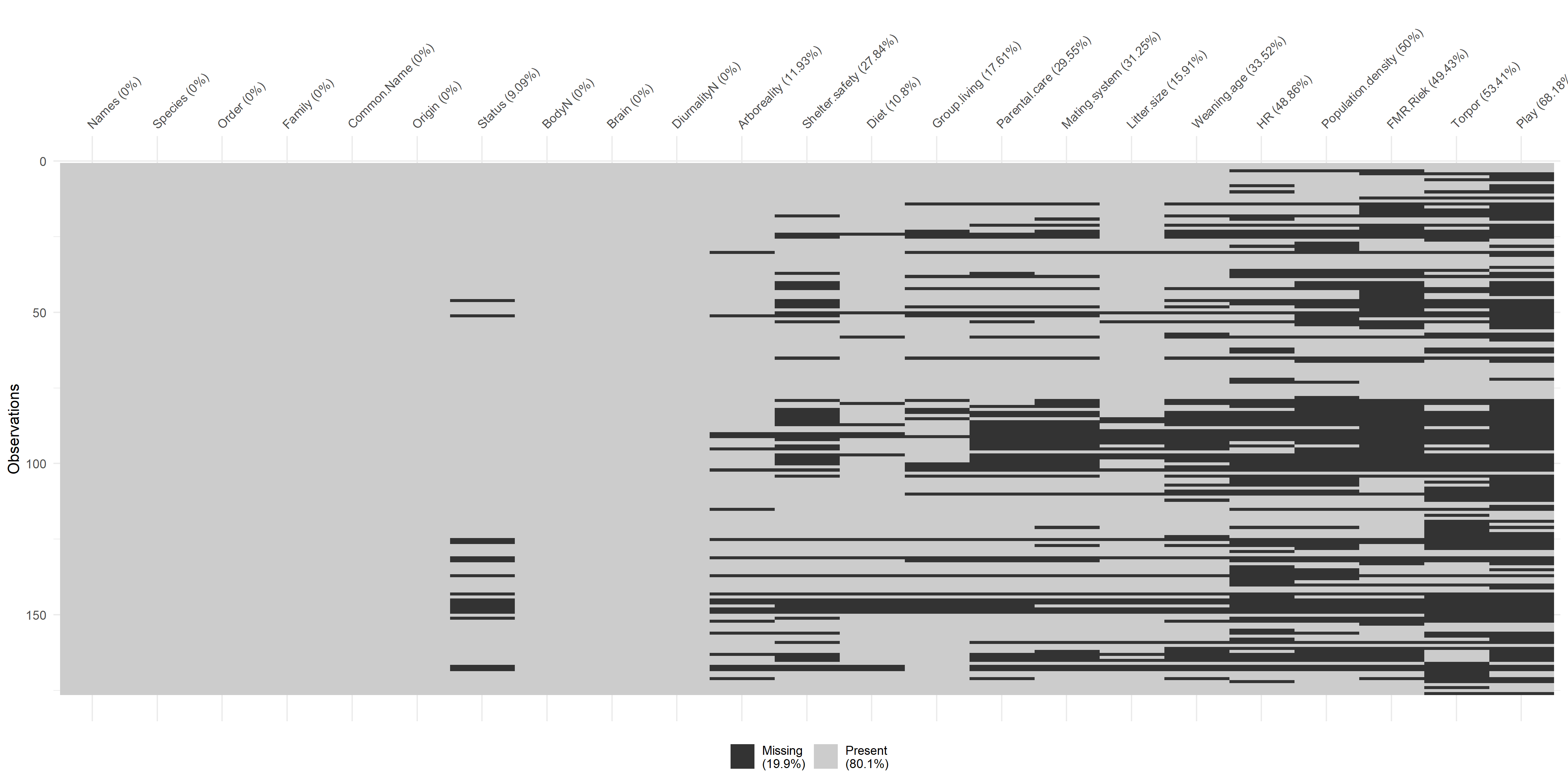

Figure 1b. Pattern of missingness and percentage of missing data per variable

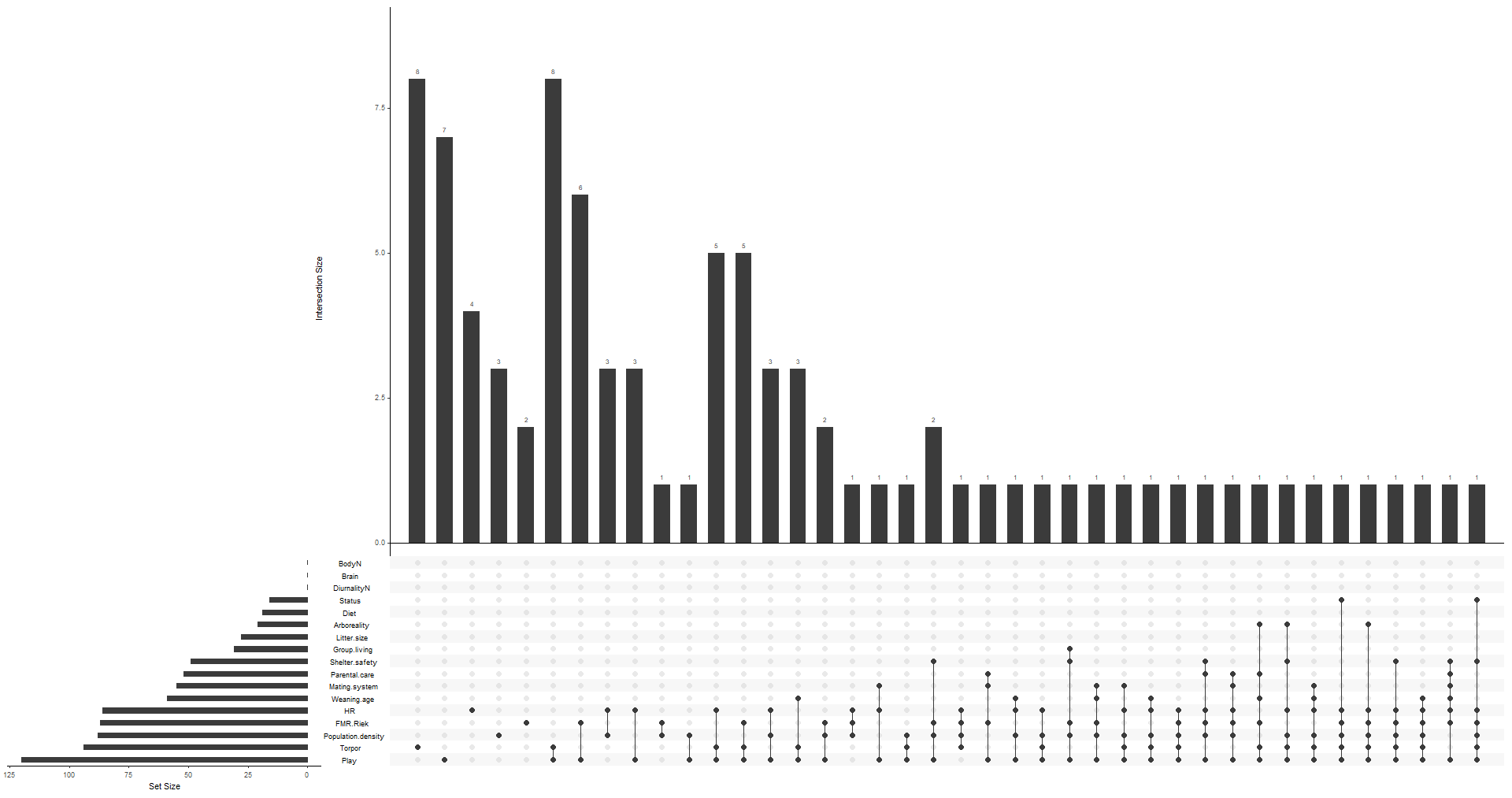

Figure 1c. NA Structure

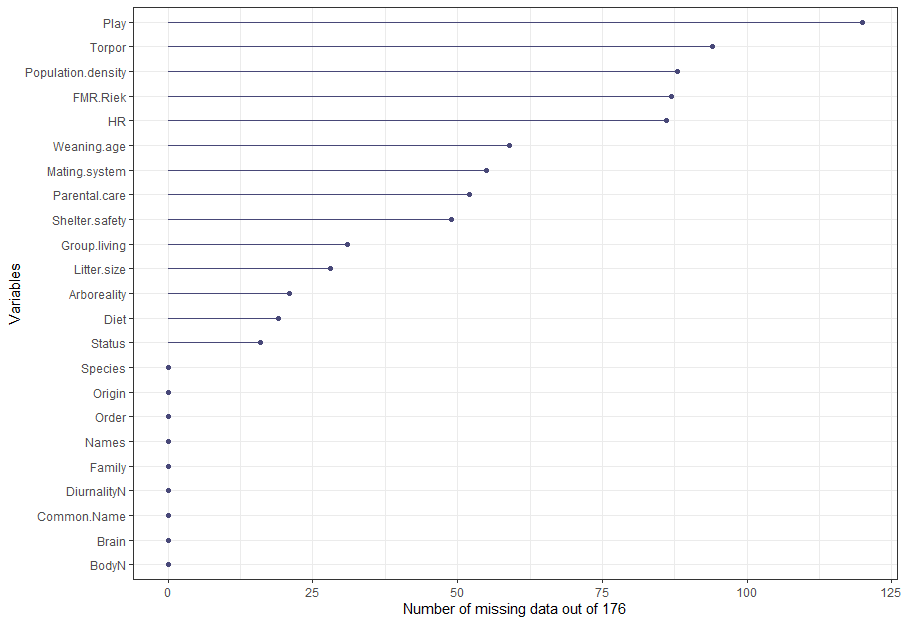

Figure 2. Number of missing data per variable out of 176 cases

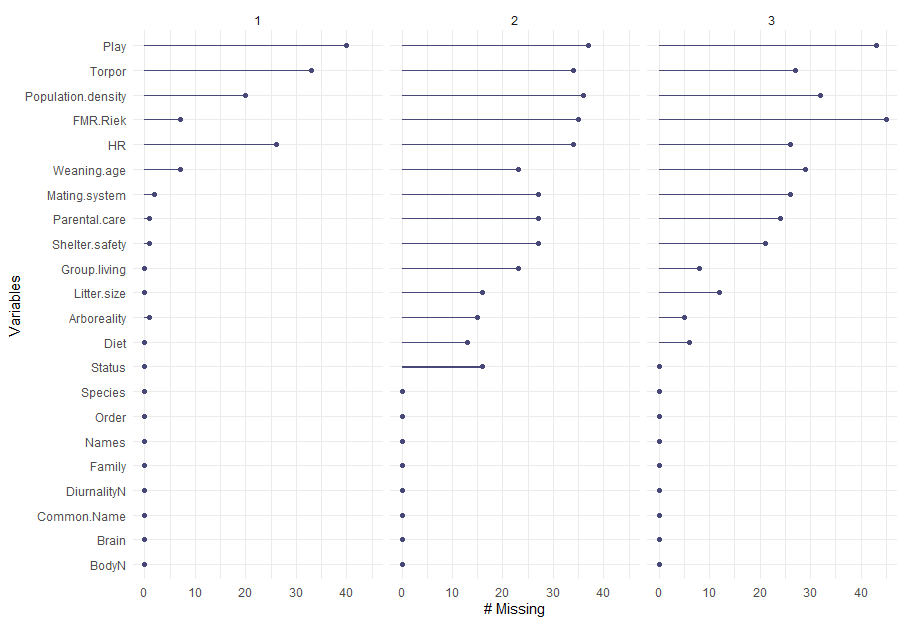

Figure 3. Number of missing data per geographic origin: 1 – Australia (n=90), 2 – New Guinea (n=40), 3 – Americas (n=45)

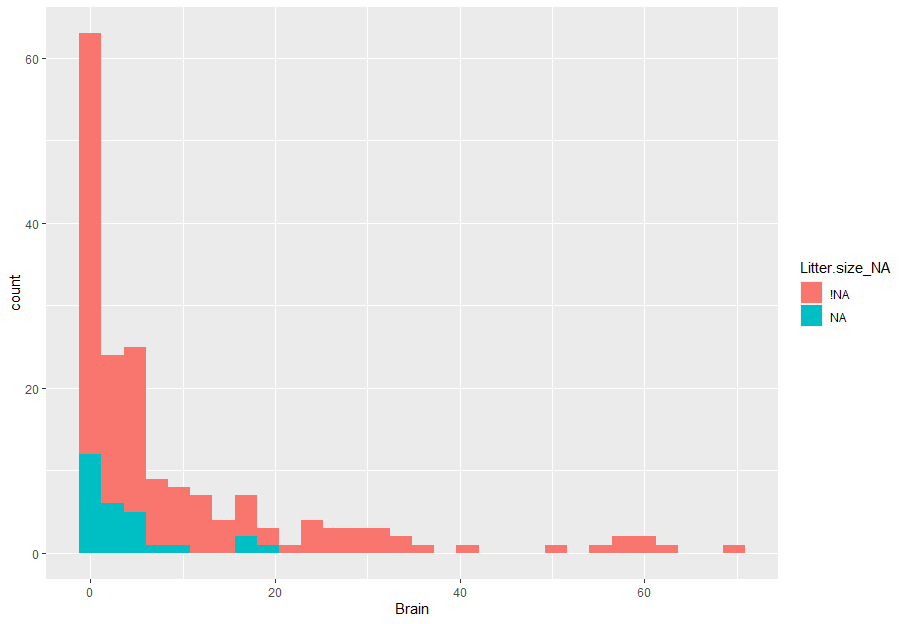

Figure 4a. Number of missing (NA) vs non-missing (!NA) data of litter size per different values of brain size. No variable showed any concerning pattern of missingness.

Phylogenetic signal in missing data

| Variable | N missing | N present | Estimated D | Probability of D random | Probability of D Brownian Motion |
| --- | --- | --- | --- | --- | --- |
| Status | 16 | 160 | 0.26 | 0 | 0.26 |
| Arboreality | 21 | 155 | 0.89 | 0.22 | 0 |
| Shelter safety | 49 | 127 | 0.52 | 0 | 0.01 |
| Diet | 19 | 157 | 0.65 | 0.01 | 0.02 |
| Group living | 31 | 145 | 0.59 | 0 | 0.01 |
| Parental care | 52 | 124 | 0.56 | 0 | 0.01 |
| Mating system | 55 | 121 | 0.45 | 0 | 0.03 |
| Litter size | 28 | 148 | 0.65 | 0 | 0 |
| Weaning age | 59 | 117 | 0.42 | 0 | 0.05 |
| Home range | 86 | 90 | 0.65 | 0 | 0 |
| Population density | 88 | 88 | 0.27 | 0 | 0.13 |
| FMR | 87 | 89 | 0.09 | 0 | 0.35 |
| Torpor | 94 | 82 | 0.66 | 0 | 0 |
| Play | 120 | 56 | 0.61 | 0 | 0 |

D is not significantly different from the Brownian motion expectation (D = 0). FMR is nearest to a random D as some of the data in it is already based on imputations. Arboreality has the highest estimated D but still does not contain phylogenetic signal.

*“D typically varies between 0 and 1. A D of 0 indicates that a trait evolves on a tree following the Brownian model (strong phylogenetic signal), and a D of 1 indicates that a trait evolves following a random model (no phylogenetic signal). D can be negative, which means that a trait evolves in a conserved way: more conserved than predicted by the Brownian model.”*

After inspecting all diagnostic plots included above, we proceed to multiple imputations.

Evaluation

Visual inspection and evaluation of ‘realistic’ imputation values of numerous species in all 25 imputed datasets was performed at random, and the imputed data seems to be realistic. Due to the nature of the missing data, many of the traits are difficult to compare with ‘realistic values’ (i.e. metabolic rate, play behavior). Due to the nature of multiple imputations, many values (including categorical variables) were different in different imputed datasets i.e. the same species was imputed as, for example, arboreal in one dataset, but as terrestrial in another.

PGLS

Table 1. Resume of PGLS models: DF – Degrees of freedom. Significant relationship between Origin 2 (New Guinea) (p=.007) and brain size, and interaction between vulnerability status 2 and body with brain size (p=.04)

| Model | Significance | DF |
| --- | --- | --- |
| Developmental | NS | 117 |
| Social | NS | 84 |
| Environmental | NS | 83 |
| Origin | Origin 2 (New Guinea)- p=.007 | 176 |
| Vulnerability status | Status2 (vulnerable):body size - p=.04 | 160 |
| Torpor | NS | 82 |
| Play | NS | 56 |
| FMR | NS | 89 |

Detailed output of the PGLS models:

DEVELOPMENTAL

Generalized least squares fit by REML

Model: log(Brain) ~ log(BodyN) + Weaning.age + Litter.size

Data: data

AIC	BIC	logLik

-8.421163 5.215776 9.210582

Correlation Structure: corBrownian

Formula: ~Names

Parameter estimate(s):

numeric(0)

Coefficients:

Value Std.Error	t-value p-value

(Intercept) -2.0337675 0.25657510 -7.926598 0.0000

log(BodyN)	0.5624441 0.01984942 28.335543 0.0000

Weaning.age 0.0000836 0.00015311 0.546064 0.5861

Litter.size -0.0281859 0.01648237 -1.710062 0.0900

Correlation:

(Intr) lg(BN) Wnng.g

log(BodyN) -0.463

Weaning.age 0.042 -0.306

Litter.size -0.393 0.282 -0.015

Standardized residuals:

Min	Q1	Med	Q3	Max

-1.82713446 -0.37804997 0.07749879 0.43441000 0.95303512

Residual standard error: 0.4491754

Degrees of freedom: 117 total; 113 residual

SOCIAL

Generalized least squares fit by REML

Model: log(Brain) ~ Group.living + Parental.care + Mating.system + log(Population.density) +	log(BodyN)

Data: data

AIC	BIC	logLik

19.85457 36.35153 -2.927284

Correlation Structure: corBrownian

Formula: ~Names

Parameter estimate(s):

numeric(0)

Coefficients:

Value Std.Error	t-value p-value

(Intercept)	-2.1401495 0.3243932 -6.597394 0.0000

Group.living	-0.0194859 0.0543611 -0.358452 0.7210

Parental.care	-0.1395289 0.1544058 -0.903650 0.3690

Mating.system	0.0207417 0.0621487 0.333744 0.7395

log(Population.density) 0.0166754 0.0174469 0.955779 0.3421

log(BodyN)	0.5911899 0.0241546 24.475225 0.0000

Correlation:

(Intr) Grp.lv Prntl. Mtng.s lg(P.)

Group.living	-0.005

Parental.care	-0.434 -0.293

Mating.system	-0.183 -0.070 -0.159

log(Population.density) -0.250 -0.210 0.019 0.340

log(BodyN)	-0.459 -0.076 0.062 0.080 0.404

Standardized residuals:

Min	Q1	Med	Q3	Max

-1.7105403 -0.3365575 0.1263874 0.5201961 1.1865854

Residual standard error: 0.4904185

Degrees of freedom: 84 total; 78 residual

ENVIRONMENTAL

Generalized least squares fit by REML

Model: log(Brain) ~ DiurnalityN + Shelter.safety + Arboreality + Diet + log(HR) + log(BodyN)

Data: data

AIC	BIC	logLik

20.57575 39.22161 -2.287874

Correlation Structure: corBrownian

Formula: ~Names

Parameter estimate(s):

numeric(0)

Coefficients:

Value Std.Error	t-value p-value

(Intercept)	-1.9449889 0.29105712 -6.682499 0.0000

DiurnalityN	0.0026436 0.02241524 0.117939 0.9064

Shelter.safety -0.0418645 0.04538671 -0.922395 0.3592

Arboreality	0.0247103 0.07168543 0.344705 0.7313

Diet	-0.0582259 0.03498454 -1.664332 0.1002

log(HR)	0.0222148 0.01171517 1.896238 0.0617

log(BodyN)	0.5535672 0.02282740 24.250124 0.0000

Correlation:

(Intr) DrnltN Shltr. Arbrlt Diet	lg(HR)

DiurnalityN	-0.063

Shelter.safety -0.067 -0.065

Arboreality	-0.325 0.049 -0.436

Diet	-0.418 -0.046 -0.034 0.207

log(HR)	0.036 0.206 0.208 -0.072 0.108

log(BodyN)	-0.381 -0.115 -0.036 -0.021 0.022 -0.400

Standardized residuals:

Min	Q1	Med	Q3	Max

-1.4385022 -0.3642219 0.2163726 0.5162295 1.1221055

Residual standard error: 0.4508372

Degrees of freedom: 83 total; 76 residual

ORIGIN

Generalized least squares fit by REML

Model: log(Brain) ~ Origin * log(BodyN)

Data: data

AIC	BIC	logLik

-33.07784 -11.12725 23.53892

Correlation Structure: corBrownian

Formula: ~Names

Parameter estimate(s):

numeric(0)

Coefficients:

Value Std.Error t-value p-value

(Intercept)	-2.2441539 0.2725458 -8.23404 0.0000

Origin2	0.3297579 0.1223180 2.69591 0.0077

Origin3	0.3332724 0.3502826 0.95144 0.3427

log(BodyN)	0.5684066 0.0169076 33.61846 0.0000

Origin2:log(BodyN) -0.0344183 0.0185696 -1.85348 0.0655

Origin3:log(BodyN) -0.0434601 0.0427487 -1.01664 0.3108

Correlation:

(Intr) Orign2 Orign3 lg(BN) O2:(BN

Origin2	-0.076

Origin3	-0.628 0.059

log(BodyN)	-0.374 0.171 0.291

Origin2:log(BodyN) 0.062 -0.947 -0.048 -0.154

Origin3:log(BodyN) 0.131 -0.068 -0.466 -0.396 0.061

Standardized residuals:

Min	Q1	Med	Q3	Max

-1.84780508 -0.42296011 -0.07007503 0.60670278 1.52166315

Residual standard error: 0.4532041

Degrees of freedom: 176 total; 170 residual

STATUS

Generalized least squares fit by REML

Model: log(Brain) ~ Status * log(BodyN)

Data: data

AIC	BIC logLik

-21.8838 -6.634519 15.9419

Correlation Structure: corBrownian

Formula: ~Names

Parameter estimate(s):

numeric(0)

Coefficients:

Value Std.Error	t-value p-value

(Intercept)	-1.9195140 0.23377689 -8.210880 0.0000

Status	-0.1425084 0.07412833 -1.922456 0.0564

log(BodyN)	0.5345605 0.02036605 26.247626 0.0000

Status:log(BodyN) 0.0224436 0.01069771 2.097979 0.0375

Correlation:

(Intr) Status lg(BN)

Status	-0.437

log(BodyN)	-0.500 0.569

Status:log(BodyN) 0.393 -0.938 -0.607

Standardized residuals:

Min	Q1	Med	Q3	Max

-1.8561615 -0.4791775 -0.1007457 0.4672986 1.5251185

Residual standard error: 0.4597327

Degrees of freedom: 160 total; 156 residual

TORPOR

Generalized least squares fit by REML

Model: log(Brain) ~ Torpor * log(BodyN)

Data: data

AIC	BIC	logLik

-17.15825 -5.374705 13.57912

Correlation Structure: corBrownian

Formula: ~Names

Parameter estimate(s):

numeric(0)

Coefficients:

Value Std.Error	t-value p-value

(Intercept)	-2.2213566 0.26033158 -8.532797 0.0000

Torpor	-0.2348034 0.22461059 -1.045380 0.2991

log(BodyN)	0.5812617 0.02597856 22.374668 0.0000

Torpor:log(BodyN) 0.0339674 0.03769176 0.901188 0.3703

Correlation:

(Intr) Torpor lg(BN)

Torpor	-0.568

log(BodyN)	-0.678 0.707

Torpor:log(BodyN) 0.464 -0.939 -0.596

Standardized residuals:

Min	Q1	Med	Q3	Max

-1.7295874 -0.5906178 0.1101043 0.4666680 1.1443593

Residual standard error: 0.3938022

Degrees of freedom: 82 total; 78 residual

PLAY

Generalized least squares fit by REML

Model: log(Brain) ~ Play * log(BodyN)

Data: data

AIC	BIC	logLik

-14.90711 -5.150887 12.45355

Correlation Structure: corBrownian

Formula: ~Names

Parameter estimate(s):

numeric(0)

Coefficients:

Value Std.Error	t-value p-value

(Intercept)	-3.0915753 0.3485394 -8.870089 0.0000

Play	0.2399924 0.1230628 1.950162 0.0566

log(BodyN)	0.6819123 0.0517292 13.182345 0.0000

Play:log(BodyN) -0.0280466 0.0199130 -1.408457 0.1649

Correlation:

(Intr) Play	lg(BN)

Play	-0.807

log(BodyN)	-0.830 0.839

Play:log(BodyN) 0.794 -0.925 -0.939

Standardized residuals:

Min	Q1	Med	Q3	Max

-1.0882984 -0.1913995 0.4231502 0.8434442 1.9350479

Residual standard error: 0.3412991

Degrees of freedom: 56 total; 52 residual

FMR

Generalized least squares fit by REML

Model: log(Brain) ~ FMR.Riek * log(BodyN)

Data: data

AIC	BIC	logLik

34.7442 46.95745 -12.3721

Correlation Structure: corBrownian

Formula: ~Names

Parameter estimate(s):

numeric(0)

Coefficients:

Value Std.Error	t-value p-value

(Intercept)	-2.3356588 0.3277660 -7.125995 0.0000

FMR.Riek	-0.0002394 0.0003741 -0.639979 0.5239

log(BodyN)	0.5897278 0.0311626 18.924188 0.0000

FMR.Riek:log(BodyN) 0.0000213 0.0000339 0.627334 0.5321

Correlation:

(Intr) FMR.Rk lg(BN)

FMR.Riek	0.111

log(BodyN)	-0.433 -0.531

FMR.Riek:log(BodyN) -0.097 -0.998 0.499

Standardized residuals:

Min	Q1	Med	Q3	Max

-1.5084653 -0.3090352 0.2075147 0.6725456 1.1756767

Residual standard error: 0.4899207

Degrees of freedom: 89 total; 85 residual
